# Integrin beta 4 promotes colorectal cancer progression by upregulating Ezrin and activating the Wnt/β-catenin signaling pathway

**DOI:** 10.1101/2024.12.19.629152

**Authors:** Jing Wang, Yi Si, Mingda Xuan, Shuangshuang Han, Kunyi Liu, Jiao Jiao, Xiaoyan Men, Hongfei Li, Jia Wang, Ting Liu, Weifang Yu

**Affiliations:** Department of Endoscopy Center, The First Hospital of Hebei Medical University, Shijiazhuang, Hebei, 050031, China; Department of Infectious Diseases, The First Hospital of Hebei Medical University, Shijiazhuang, Hebei, 050031, China

**Author notes:** **Lead Corresponding Author: Weifang Yu, M.D.,** Professor; Department of Endoscopy Center, The First Hospital of Hebei Medical University, No. 89 Donggang Road, Shijiazhuang, Hebei, 050031, China. **Co-Corresponding Author: Ting Liu, M.D.;** Department of Endoscopy Center, The First Hospital of Hebei Medical University, No. 89 Donggang Road, Shijiazhuang, Hebei, 050031, China. These authors contributed equally to this work (Jing Wang, Yi Si, Mingda Xuan).

**Keywords:** colorectal cancer, integrin beta 4, Ezrin, Wnt/β-catenin signaling, metastasis, prognostic biomarker, therapeutic target

## Abstract

**Objectives:** Colorectal cancer (CRC) is a major cause of cancer-related mortality worldwide. Integrin beta 4 (ITGB4) has been previously identified as being overexpressed in CRC; however, its precise oncogenic mechanism remains unclear. The present study aimed to elucidate the functional role of ITGB4 in CRC progression and identify its downstream molecular effectors to provide new insights for targeted therapy.

**Methods:** The biological functions of ITGB4 were investigated in CRC cell lines (SW480 and HCT116) using a series of *in vitro* assays, including CCK-8, colony formation, Transwell migration and invasion, and flow cytometry for apoptosis following ITGB4 knockdown. An *in vivo* xenograft mouse model was used to evaluate the effect of ITGB4 on tumor growth. Downstream targets were screened using RNA sequencing (RNA-seq) and validated by co-immunoprecipitation and co-immunofluorescence. The underlying signaling pathway was investigated by Western blotting and functional rescue experiments.

**Results:** Knockdown of ITGB4 significantly suppressed CRC cell proliferation, migration, and invasion, while promoting apoptosis *in vitro*. Similarly, silencing ITGB4 markedly inhibited tumor growth in the *in vivo* xenograft model. RNA-seq analysis identified Ezrin (EZR) as a key downstream target of ITGB4, and a direct protein-protein interaction was confirmed between them. Mechanistically, ITGB4 knockdown decreased the expression of EZR at both the mRNA and protein levels. ITGB4 was demonstrated to exert its pro-tumorigenic effects through the regulation of EZR, which subsequently activated the Wnt/β-catenin signaling pathway. Interestingly, EZR overexpression partially restored ITGB4 levels, suggesting a potential positive feedback loop via Wnt/β-catenin signaling that further amplifies this oncogenic axis. Notably, the malignant phenotypes suppressed by ITGB4 silencing were significantly rescued by the overexpression of EZR.

**Conclusion:** The present study identified a novel ITGB4/EZR/Wnt/β-catenin signaling axis in colorectal cancer. ITGB4 promotes CRC progression by modulating EZR expression and subsequently activating the Wnt/β-catenin pathway. These findings highlight ITGB4 as a potential prognostic biomarker and a promising therapeutic target for CRC.

## Introduction

Metastasis remains the primary cause of mortality in colorectal cancer (CRC) patients [1]. The metastatic cascade is a complex, multi-step process initiated by the detachment of cancer cells from the primary tumor, a step heavily regulated by cell-matrix adhesion molecules [2]. Alterations in the expression of integrins and their interactions with the extracellular matrix (ECM) are fundamental to this process, allowing tumor cells to acquire invasive properties [3]. Consequently, there is an urgent need to identify novel therapeutic targets that can inhibit the proliferation, invasion, and migration of CRC cells. The advent of high-throughput technologies and public biological databases has accelerated research into the genetic and epigenetic alterations driving CRC, revealing that specific gene mutations are pivotal to its pathogenesis [4]. In-depth investigation of these aberrantly expressed genes is crucial for understanding CRC biology and developing effective strategies for early diagnosis and precision treatment.

Our research group previously identified Integrin Beta 4 (ITGB4) as a gene of significant diagnostic and therapeutic potential in CRC, leveraging advanced technologies such as mass cytometry (CyTOF), gene chips, and protein arrays. This discovery led to a national invention patent (Patent Number: ZL 2017 1 1148747.1) [5]. ITGB4, a member of the integrin family, is predominantly expressed on the basal surface of epithelial cells and forms hemidesmosomes by dimerizing with integrin alpha 6 (ITGA6) [6]. Accumulating evidence indicates that ITGB4 is overexpressed in various malignancies, including breast, bladder, cervical, head and neck, lung, and pancreatic cancers, where its expression often correlates with tumor invasion and poor prognosis [7–14]. Our prior studies corroborated these findings in CRC, demonstrating that elevated ITGB4 levels in both tissues and serum are associated with adverse clinicopathological features and reduced overall survival [15, 16]. Gene co-expression analysis suggested that ITGB4’s regulatory role in CRC involves multiple signaling pathways related to cell growth, migration, and apoptosis. However, the specific molecular mechanism through which ITGB4 functions in CRC has not been fully elucidated.

Integrins relay signals by connecting the ECM to the intracellular actin cytoskeleton. This linkage is often mediated by scaffolding proteins such as the Ezrin-Radixin-Moesin (ERM) family, which crosslink actin filaments to plasma membrane proteins [17]. Given this functional synergy, we hypothesized that ITGB4 might orchestrate CRC progression through specific interactions with cytoskeletal linkers like Ezrin. This study aimed to systematically investigate the role of ITGB4 in the initiation and progression of CRC. We first validated its oncogenic functions in vitro and in vivo. Subsequently, through RNA sequencing (RNA-seq) and a multi-step bioinformatic filtering process, we identified Ezrin (EZR) as a novel downstream target of ITGB4. We demonstrate that ITGB4 interacts with and positively regulates the expression of EZR, thereby activating the Wnt/β-catenin signaling pathway to promote CRC progression. Our findings delineate the ITGB4/EZR/Wnt/β-catenin axis as a critical regulatory network in CRC, offering a potential new target for precision therapeutic intervention.

## Materials and Methods

### Cell Culture

The human colorectal cancer cell lines SW480 (ATCC® CCL-228™) and HCT116 (ATCC® CCL-247™) were purchased from the American Type Culture Collection (ATCC, Manassas, VA, USA). Cells were cultured in Dulbecco’s Modified Eagle’s Medium (DMEM; Gibco, Grand Island, NY, USA) supplemented with 10% fetal bovine serum (FBS; Gibco) and 1% penicillin-streptomycin. Cultures were maintained in a humidified incubator at 37°C with 5% CO_2_. All cells used in experiments were in the logarithmic growth phase. Cell line authentication was performed prior to the study, and routine testing confirmed the absence of mycoplasma contamination.

### siRNA, shRNA, Plasmids, and Transfections

For transient knockdown, small interfering RNAs (siRNAs) targeting ITGB4 (siITGB4) and a negative control siRNA (siNC) were synthesized by GenePharma Co., Ltd. (Shanghai, China). For EZR overexpression, a human EZR open reading frame was cloned into the pcDNA3.1 vector (pcDNA3.1-EZR), with the empty vector serving as a control (pcDNA3.1-vector); these were also obtained from GenePharma. For stable knockdown, short hairpin RNAs targeting ITGB4 (shITGB4) and a non-targeting control (shNC) were cloned into a lentiviral vector by GenePharma. The sequences for all siRNAs and shRNAs used are listed in Supplementary Table 1. Transfections were performed using Lipofectamine 3000 (Invitrogen, Carlsbad, CA, USA) according to the manufacturer’s protocol when cell confluence reached 80-90%.

### Generation of Stable Cell Lines

To establish stable ITGB4-knockdown cell lines, HCT116 cells were transfected with shITGB4 or shNC vectors. Forty-eight hours post-transfection, the cells were subjected to selection with 600 µg/ml G418 (Neomycin). The selection medium was refreshed every 2-3 days. After approximately one week, when significant cell death was observed, the G418 concentration was reduced to 300 µg/ml for maintenance. After 10-14 days of selection, resistant cell colonies were pooled, expanded, and cultured without G418. The efficiency of stable knockdown was confirmed by qRT-PCR and Western blot analysis.

### RNA Extraction and Quantitative Real-Time PCR (qRT-PCR)

Total RNA was extracted from cells using the RNA-Easy Kit (Vazyme Biotech Co., Ltd., Nanjing, China). cDNA was synthesized from 1 µg of total RNA using the PrimeScript RT Reagent Kit (Takara, Beijing, China). qRT-PCR was performed on a 4800 Real-Time PCR System using SYBR Green Master Mix (Vazyme). The thermal cycling conditions were: 95°C for 5 min, followed by 40 cycles of 95°C for 10 s and 60°C for 30 s. Relative gene expression was calculated using the 2^−ΔΔCt^ method [18], with β-actin serving as the internal control. Primer sequences are listed in Supplementary Table 2.

### Western Blot Analysis

Cells were lysed in RIPA buffer (Solarbio, Beijing, China) supplemented with a protease inhibitor cocktail (PMSF; Solarbio). Protein concentrations were determined using a BCA protein assay kit (Solarbio). Equal amounts of protein (30 μg) were separated by 10% SDS-PAGE and transferred to polyvinylidene fluoride (PVDF) membranes (Merck Millipore, Billerica, MA, USA). The membranes were blocked with 5% non-fat milk in TBST for 1 h at room temperature and then incubated overnight at 4°C with primary antibodies. The primary antibodies used were: anti-ITGB4 (1:1000, Abcam, ab261778), anti-EZR (1:1000, Proteintech, 26056-1-AP), anti-β-Catenin (1:1000, Proteintech, 51067-2-AP), anti-c-Myc (1:1000, Proteintech, 10828-1-AP), anti-Cyclin D1 (1:500, Proteintech, 60186-1-Ig), and anti-β-actin (1:1500, ZSBG-Bio, TA-09). After washing, membranes were incubated with corresponding HRP-conjugated secondary antibodies. Immunoreactive bands were visualized using an Odyssey scanning system (LI-COR Biosciences, Lincoln, NE, USA). Band intensities were quantified using ImageJ software (National Institutes of Health, Bethesda, MD, USA).

### Cell Proliferation Assays

For the Cell Counting Kit-8 (CCK-8) assay, cells were seeded into 96-well plates at a density of 1×10^3^ cells/well. At 24, 48, 72, and 96 h, 10 µl of CCK-8 reagent (Dojindo, Tokyo, Japan) was added to each well, followed by a 2 h incubation at 37°C. The absorbance at 450 nm was measured using a microplate reader (Promega, Madison, WI, USA). For the colony formation assay, cells were seeded into 6-well plates at 500 cells/well and cultured for 10-14 days, with the medium changed every 3 days. Colonies were fixed with 4% paraformaldehyde, stained with 0.1% crystal violet, and colonies containing >50 cells were counted.

### Cell Migration and Invasion Assays

For the wound healing assay, cells were grown to 100% confluence in 6-well plates. A scratch was made using a 200 µl pipette tip. After washing with PBS, cells were cultured in serum-free medium. Images were captured at 0 and 48 h. The migration rate was calculated based on the change in wound width. For Transwell assays, 1×10^5^ cells in 200 µl of serum-free medium were seeded into the upper chamber of a Transwell insert (8.0 μm pore size; Corning, NY, USA). The lower chamber contained 700 µl of DMEM with 10% FBS. For the invasion assay, the upper chamber was pre-coated with Matrigel (BD Biosciences, San Jose, CA, USA). After 48 h of incubation, non-migrated/invaded cells were removed from the upper surface. Cells on the lower surface were fixed with 4% paraformaldehyde, stained with 0.1% crystal violet, and counted in five random fields under a microscope.

### Apoptosis Assay

Cell apoptosis was assessed using the Annexin V-FITC/PI Apoptosis Detection Kit (NeoBioscience, Shenzhen, China). After 48 h of transfection, cells were harvested, washed with PBS, and resuspended in binding buffer. Cells were then stained with Annexin V-FITC and Propidium Iodide (PI) for 15 min in the dark. The percentage of apoptotic cells was determined by flow cytometry (BD Biosciences) and analyzed using FlowJo software.

### Animal Studies

A total of 12 five-week-old male BALB/c nude mice, weighing 18-22g, were housed in a specific pathogen-free (SPF) facility under a 12-hour light/dark cycle with free access to food and water. They were randomly divided into two groups (n=6 per group). HCT116 cells (2×10^6^) stably expressing shNC or shITGB4 were suspended in 100 µl of serum-free DMEM and injected subcutaneously into the right flank of each mouse. Tumor growth was monitored every 3 days by measuring the length (L) and width (W) with calipers. Tumor volume was calculated using the formula: V = (L × W^2^) / 2. After 21 days, all animals were euthanized by cervical dislocation after anesthesia, and tumors were excised, weighed, and processed for histological analysis. Tumors were fixed in 4% paraformaldehyde for subsequent hematoxylin and eosin (HE) and immunohistochemistry (IHC) staining. The study protocols were approved by the Experimental Animal Care and Use Committee and Ethics Committee of The First Hospital of Hebei Medical University (Approval No. 20220395).

### RNA Sequencing (RNA-seq) and Bioinformatic Analysis

Total RNA was extracted from SW480 cells transfected with siNC or siITGB4 (three biological replicates per group). Library construction and sequencing were performed by Novogene Technology Co., Ltd. (Beijing, China) on an Illumina NovaSeq 6000 platform. After quality control, clean reads were aligned to the human reference genome (GRCh38). Differential expression analysis was performed using DESeq2. Genes with an adjusted p-value < 0.05 and a log2|Fold Change| > 1.0 were considered differentially expressed genes (DEGs).

### Immunohistochemistry (IHC)

Paraffin-embedded tumor sections were deparaffinized, rehydrated, and subjected to antigen retrieval. Sections were then incubated with an anti-Ki67 antibody (1:100; Proteintech, 27309-1-AP) overnight at 4°C. Staining was performed using a two-step detection kit (ZSBG-Bio, Beijing, China) and visualized with DAB. Images were captured, and the staining intensity was quantified as the average optical density (AOD) using ImageJ software.

### Co-immunoprecipitation (Co-IP) and Co-immunofluorescence (Co-IF)

For Co-IP, SW480 cell lysates were incubated with anti-ITGB4 antibody (10 μg; Proteintech), anti-EZR antibody (10 μg; Proteintech), or control IgG (10 μg; Proteintech) using the Pierce™ Classic Magnetic IP/Co-IP Kit (Thermo Scientific, 88804) according to the manufacturer’s instructions. Immunoprecipitated proteins were analyzed by Western blotting. For Co-IF, SW480 cells grown on glass coverslips were fixed with 4% paraformaldehyde, permeabilized with 0.2% Triton X-100, and blocked with 2% BSA. Cells were then incubated with primary antibodies against ITGB4 (1:50, Santa Cruz, sc-13543) and EZR (1:50, Proteintech, 26056-1-AP) overnight at 4°C. After washing, cells were incubated with Cy3-conjugated anti-rabbit and FITC-conjugated anti-mouse secondary antibodies. Nuclei were counterstained with DAPI. Images were acquired using a Zeiss laser scanning confocal microscope (Oberkochen, Germany).

### Public Database Analysis

ITGB4 expression data in CRC and adjacent normal tissues were obtained from The Cancer Genome Atlas (TCGA) and Gene Expression Profiling Interactive Analysis 2 (GEPIA2; http://gepia2.cancer-pku.cn/) databases. We prioritized the TCGA and GEPIA2 datasets due to their large sample sizes, standardized high-throughput sequencing platforms, and comprehensive clinical annotations, which provide robust statistical power for differential expression and survival analyses compared to smaller, individual datasets. Inclusion criteria involved datasets containing paired tumor and adjacent normal tissues with complete follow-up information. Although clinicopathological variables such as tumor stage and subtype are available in these datasets, the current analysis focused on overall differential expression and survival correlations to identify potential targets, without stratification by specific parameters. The prognostic value of gene expression was assessed using the Kaplan-Meier plotter (https://kmplot.com/analysis/). Subcellular localization data were retrieved from GeneCards (https://www.genecards.org/).

### Statistical Analysis

All experiments were performed with at least three independent replicates. Data are presented as the mean ± standard deviation (SD). Statistical analyses were performed using GraphPad Prism 8.0 and SPSS 21.0. Differences between two groups were analyzed using a two-tailed Student’s t-test. Comparisons among multiple groups were performed using one-way or two-way ANOVA followed by an appropriate post-hoc test. A p-value < 0.05 was considered statistically significant.

## Results

### ITGB4 is upregulated in colorectal cancer and correlates with poor prognosis

To corroborate our previous findings, we first analyzed ITGB4 expression using public databases. Analysis of TCGA and GEPIA2 datasets confirmed that ITGB4 mRNA expression was significantly higher in CRC tissues compared to adjacent normal tissues (**Figure 1A-C**). Furthermore, Kaplan-Meier survival analysis revealed that patients with high ITGB4 expression had significantly poorer overall survival (OS) than those with low expression (**Figure 1D**). These results are consistent with our prior work [6] and collectively underscore that ITGB4 is an oncogenic factor in CRC, and its overexpression is associated with unfavorable patient outcomes, highlighting its potential as a prognostic biomarker.

**Figure 1.**
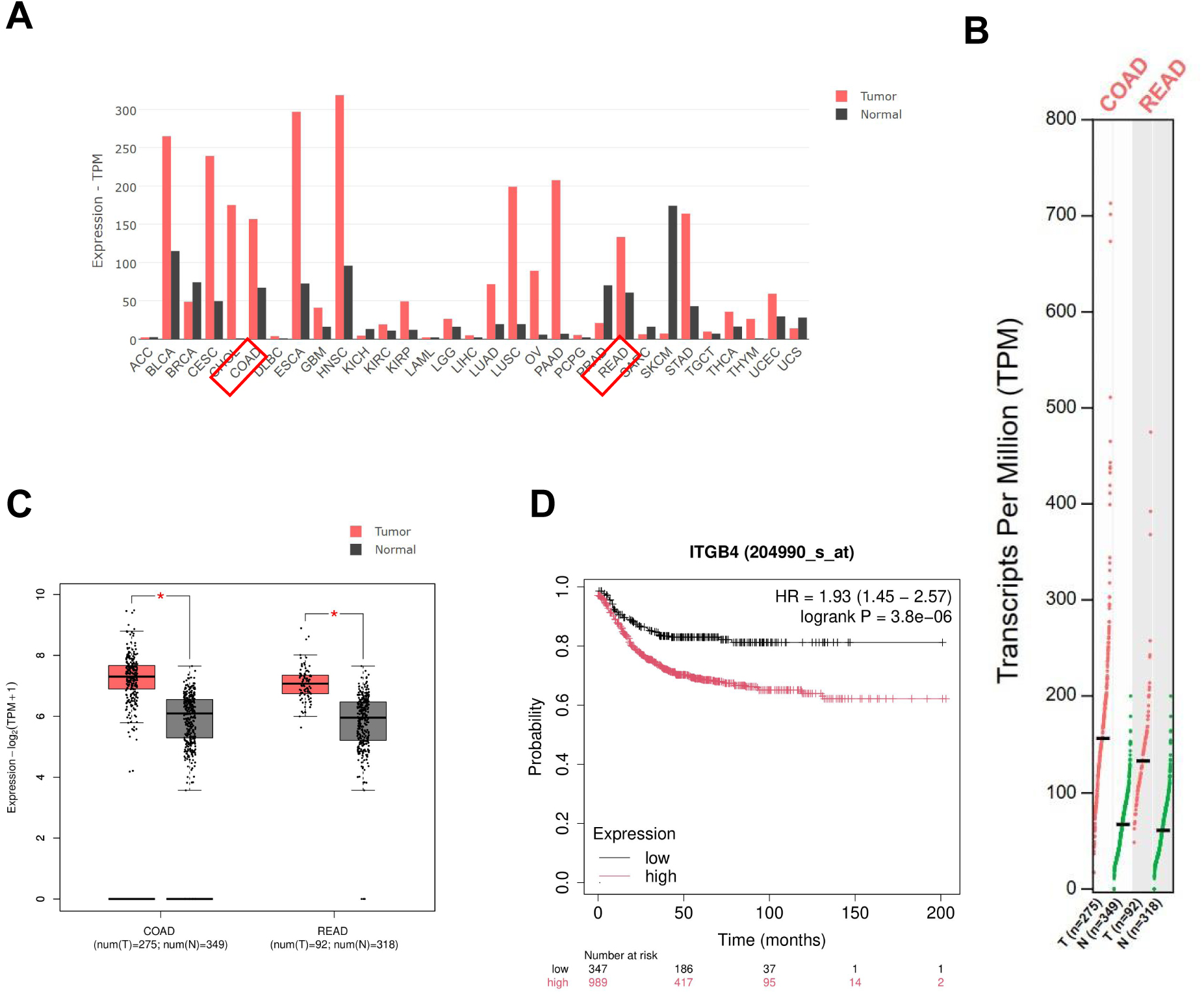
ITGB4 is upregulated in colorectal cancer and associated with poor prognosis. (A-C) Analysis of ITGB4 mRNA expression in colorectal cancer tissues and adjacent normal tissues using the TCGA and GEPIA2 databases. (D) Kaplan-Meier analysis demonstrating the correlation between ITGB4 expression levels and overall survival in colorectal cancer patients.

### Downregulation of ITGB4 suppresses malignant phenotypes of colorectal cancer cells in vitro

To investigate the biological function of ITGB4 in CRC, we first silenced its expression in SW480 and HCT116 cells using siRNAs. Four different siRNA sequences were tested, and the siITGB4-681 sequence, which demonstrated the most potent knockdown efficiency at both mRNA and protein levels, was selected for subsequent experiments (**Figure 2A, B**). Transient transfection with siITGB4-681 effectively reduced ITGB4 mRNA and protein expression in both cell lines (**Figure 2C-F**). Functional assays revealed that ITGB4 knockdown significantly inhibited cell proliferation, as determined by CCK-8 assays (**Figure 2G, H**), and suppressed clonogenic ability, as shown by colony formation assays (**Figure 2I, J**). Moreover, flow cytometry analysis indicated that ITGB4 silencing significantly increased the proportion of apoptotic cells (**Figure 2K, L**). Transwell assays demonstrated a marked reduction in both the migratory and invasive capacities of CRC cells upon ITGB4 knockdown (**Figure 2M, N**). Together, these in vitro results suggest that ITGB4 plays a critical role in promoting CRC cell proliferation, survival, migration, and invasion.

**Figure 2.**
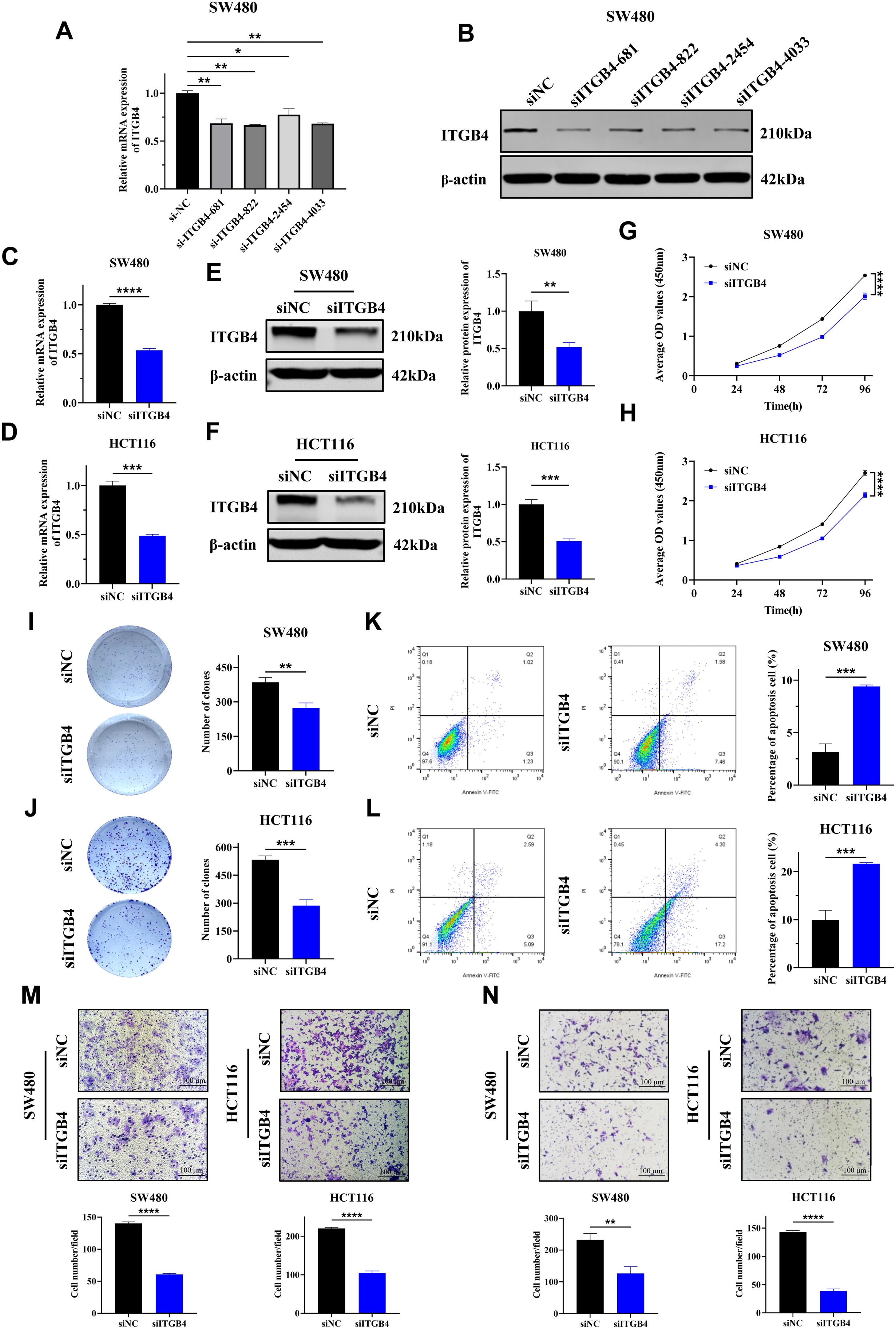
Knockdown of ITGB4 inhibits proliferation, migration, and invasion and promotes apoptosis in colorectal cancer cells. (A) qRT-PCR validation of knockdown efficiency for four different siITGB4 sequences in SW480 cells. (B) Western blot validation of knockdown efficiency for the four siITGB4 sequences. (C, D) Verification of siITGB4 knockdown efficiency at the mRNA level in SW480 and HCT116 cells via qRT-PCR. (E, F) Verification of siITGB4 knockdown efficiency at the protein level via Western blot. (G, H) CCK-8 assays showing the effect of ITGB4 knockdown on the proliferation of SW480 and HCT116 cells. (I, J) Colony formation assays demonstrating the effect of ITGB4 knockdown on the clonogenic ability of CRC cells. (K, L) Flow cytometry analysis showing the effect of ITGB4 knockdown on apoptosis. (M, N) Transwell assays showing the effect of ITGB4 knockdown on the migration and invasion abilities of SW480 and HCT116 cells (Scale bar = 100 μm). Data are presented as mean ± SD. *P < 0.05, **P < 0.01, ***P < 0.001.

### Knockdown of ITGB4 inhibits colorectal cancer growth in vivo

To extend our in vitro findings, we assessed the role of ITGB4 in tumor growth using a xenograft model. We established an HCT116 cell line with stable ITGB4 knockdown (shITGB4), confirming sustained suppression of ITGB4 expression (**Figure 3A, B**). These cells, along with control cells (shNC), were subcutaneously injected into nude mice. Tumor growth was monitored over 21 days. The tumors formed by shITGB4 cells grew significantly slower and were substantially smaller and lighter at the end of the experiment compared to tumors from the shNC group (**Figure 3C-F**). The reduced ITGB4 expression in the xenograft tumors was verified by qRT-PCR and Western blot (**Figure 3G, H**). HE staining confirmed the CRC tissue morphology in the tumors (**Figure 3I**). Furthermore, IHC staining for the proliferation marker Ki-67 showed significantly lower positivity in the shITGB4 group, indicating reduced cell proliferation (**Figure 3J, K**). These in vivo data provide strong evidence that ITGB4 is essential for promoting CRC tumorigenesis.

**Figure 3.**
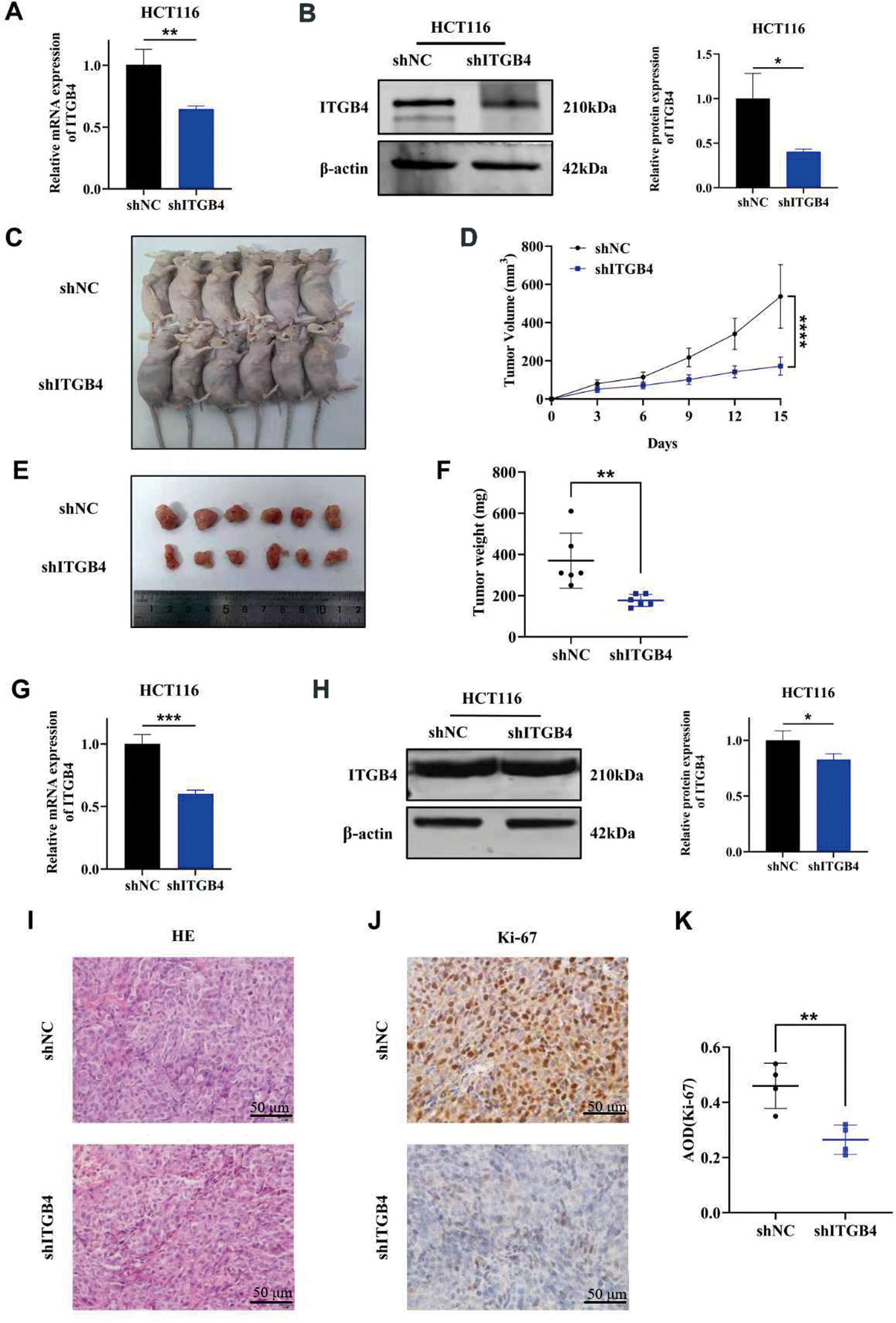
ITGB4 silencing suppresses colorectal cancer growth in vivo. (A, B) qRT-PCR and Western blot analysis verifying the stable knockdown efficiency of shITGB4 in HCT116 cells. (C) Representative images of tumor-bearing nude mice from the shNC and shITGB4 groups. (D) Tumor growth curves for both groups over 21 days. (E, F) Images and weights of the excised xenograft tumors. (G, H) Verification of ITGB4 knockdown in xenograft tumor tissues. (I) Representative hematoxylin and eosin (HE) staining of tumor sections. Scale bars = 50 μm. (J, K) Representative immunohistochemical (IHC) staining and quantification of Ki-67 in tumor sections. Scale bars = 50 μm. Data are presented as mean ± SD. **P < 0.01, ****P < 0.0001.

### Identification of EZR as a downstream target of ITGB4 in colorectal cancer

To uncover the molecular mechanisms downstream of ITGB4, we performed RNA-seq on SW480 cells following ITGB4 knockdown. This analysis identified 572 DEGs, of which 296 were downregulated and 276 were upregulated (**Figure 4A, B**). To pinpoint key downstream effectors, we devised a stringent, multi-step filtering strategy. First, we focused on the 296 downregulated genes from our RNA-seq data. This list was then intersected with two externally derived gene sets: (1) genes known to be co-expressed with ITGB4 in CRC patient cohorts (from the GEPIA database) and (2) genes that are significantly upregulated in CRC tissues compared to normal tissues (from the GEPIA2 database). This intersection yielded a refined list of 35 candidate genes. To further narrow the selection, we prioritized genes whose protein products share a subcellular localization with ITGB4 (i.e., plasma membrane or cytoplasm, based on GeneCards data). This criterion was selected because ITGB4 is a transmembrane receptor; therefore, its immediate downstream signal transducers are likely to be physically proximal, residing at the membrane or in the sub-membrane cortical cytoplasm. This systematic process identified five high-confidence candidate target genes: ITGA6, CXADR, EZR, GPRC5C, and EPHA2 (**Figure 4C**). We validated the regulatory relationship by confirming that knockdown of ITGB4 led to a significant decrease in the mRNA levels of all five candidate genes in both SW480 and HCT116 cells (**Figure 4D, E**). Among the candidates, EZR, CXADR, and ITGA6 showed predominantly plasma membrane localization, similar to ITGB4, whereas GPRC5C showed mixed localization. This spatial proximity is a prerequisite for direct protein-protein interaction. Among these candidates, EZR was selected for further investigation based on its strong positive correlation with ITGB4 expression in CRC patient samples (**Figure 5C**), its consistent upregulation in CRC tissues (**Figure 5B**), its shared subcellular localization (**Figure 5A**), and its strong association with poor patient prognosis (**Figure 5D**), which mirrored the prognostic significance of ITGB4.

**Figure 4.**
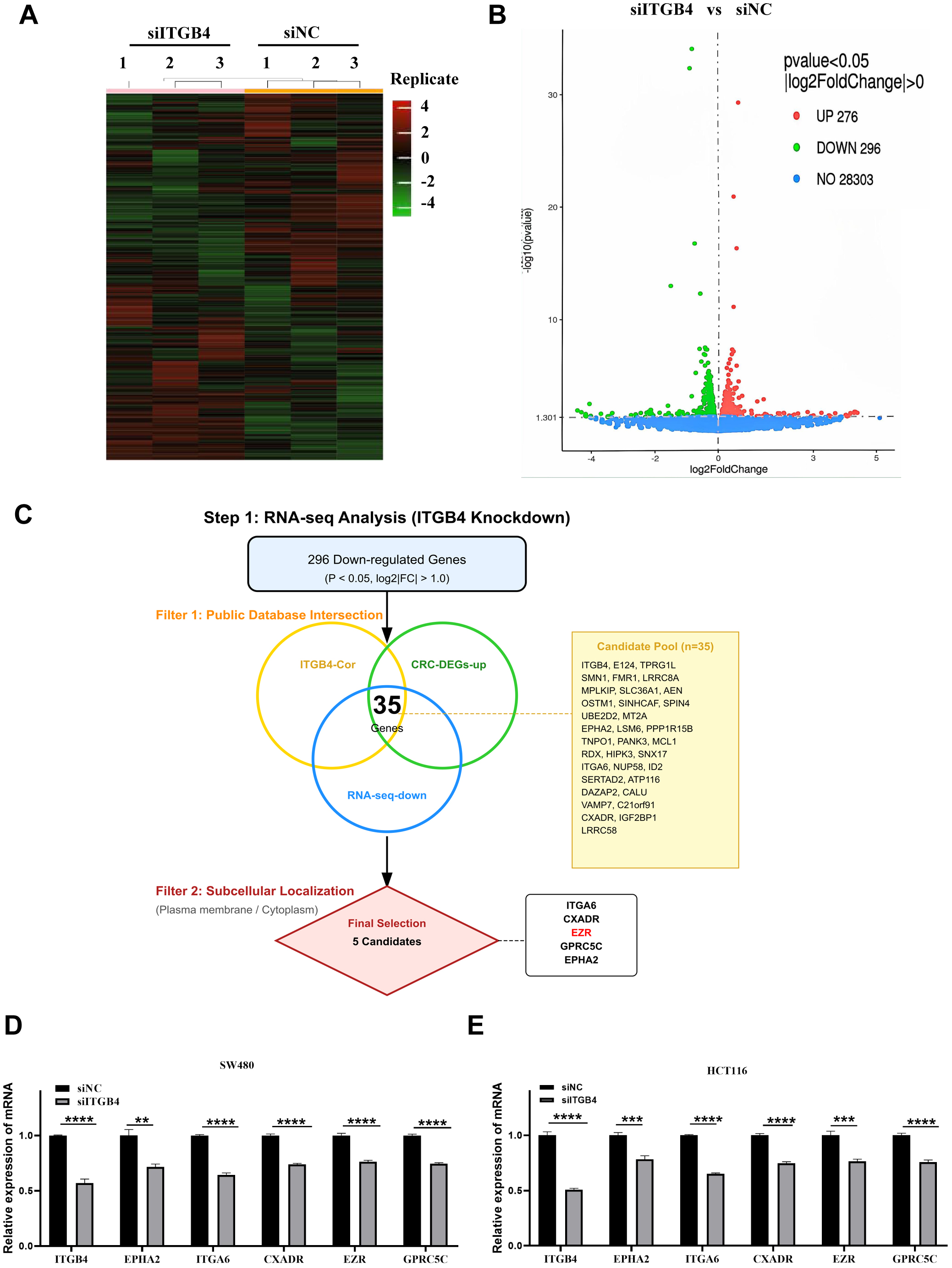
Systematic screening identifies candidate downstream targets of ITGB4. (A) Heatmap of differentially expressed genes (DEGs) identified by RNA-seq analysis of SW480 cells after ITGB4 knockdown. (B) Volcano plot illustrating the distribution of DEGs. (C) Schematic flowchart illustrating the multi-step filtering strategy. A Venn diagram was used to intersect ITGB4-downregulated genes (from RNA-seq) with genes co-expressed with ITGB4 and upregulated in CRC (from public databases), identifying 35 candidate genes. These candidates were further filtered by subcellular localization to identify the final five targets. (D, E) qRT-PCR validation of the mRNA levels of the five candidate target genes following ITGB4 knockdown in SW480 and HCT116 cells.

**Figure 5.**
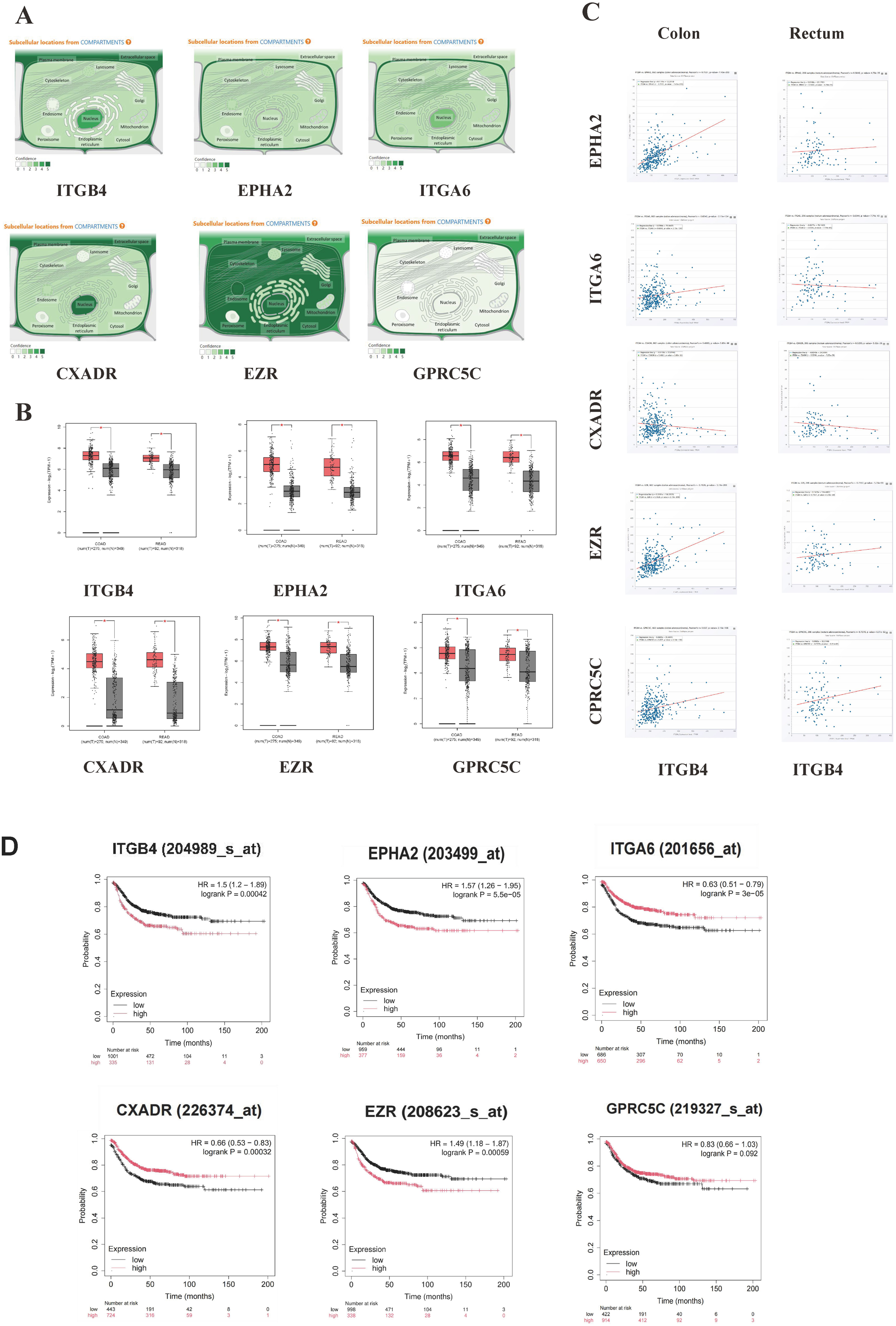
Bioinformatic analysis highlights EZR as a prime candidate downstream of ITGB4. (A) Subcellular localization of ITGB4 and the five candidate genes as annotated in the GeneCards database. (B) Expression analysis of the five candidate genes in CRC versus adjacent normal tissues using the GEPIA2 database. (C) Correlation analysis of ITGB4 expression with each of the five candidate genes in the TCGA-COAD cohort. (D) Kaplan-Meier survival analysis showing the prognostic significance of the five candidate genes in colorectal cancer patients.

### ITGB4 interacts with and positively regulates the expression of EZR

Given the strong correlational evidence, we next investigated a potential physical interaction between ITGB4 and EZR. Co-IP assays in SW480 cells demonstrated that endogenous ITGB4 could be immunoprecipitated with an anti-EZR antibody, and conversely, EZR was pulled down with an anti-ITGB4 antibody, confirming that the two proteins exist in the same complex (**Figure 6A**). Co-IF staining further revealed that ITGB4 and EZR extensively co-localize at the plasma membrane of CRC cells (**Figure 6B**). This interaction is also supported by protein-protein interaction network analysis from the STRING database (**Figure 6C**). We then confirmed that ITGB4 regulates EZR expression. As shown in **Figure 6F and G**, knockdown of ITGB4 in SW480 cells resulted in a significant reduction of both EZR mRNA and protein levels. To assess the functional hierarchy, we performed rescue experiments. After establishing an efficient EZR overexpression system (**Figure 6D, E**), we co-transfected cells with siITGB4 and the EZR overexpression plasmid. Interestingly, overexpression of EZR partially restored the suppressed levels of ITGB4 mRNA and protein induced by siITGB4 (**Figure 6H, I**). Given that EZR functions as a downstream effector that activates Wnt/β-catenin signaling, this concomitant increase in ITGB4 suggests the existence of a positive feedback loop, wherein hyperactive Wnt/β-catenin signaling transcriptionally upregulates ITGB4 to further sustain the malignant progression. Collectively, these data establish that ITGB4 interacts with EZR and positively regulates its expression at both transcriptional and post-transcriptional levels.

**Figure 6.**
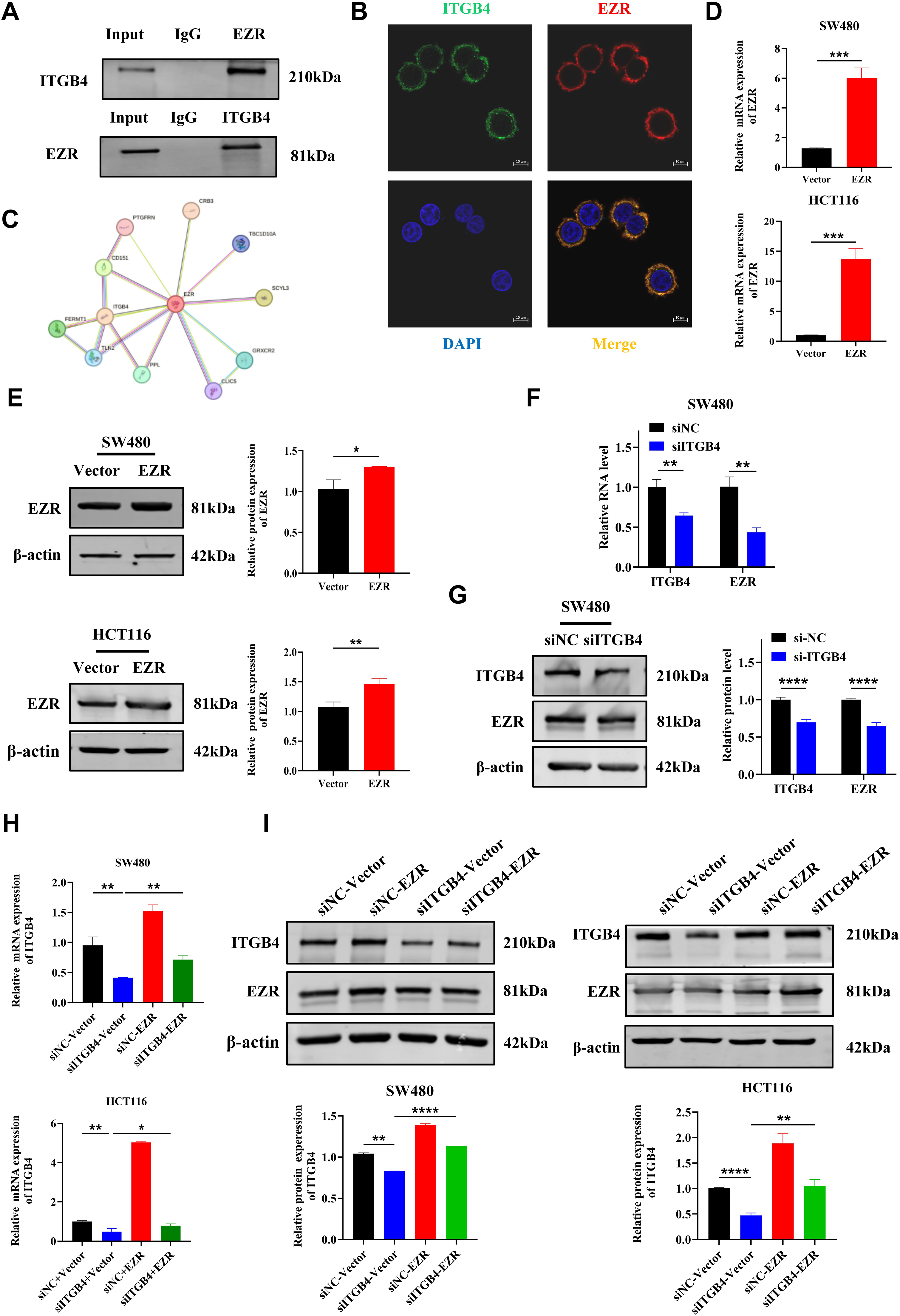
ITGB4 interacts with EZR and regulates its expression. (A) Co-immunoprecipitation (Co-IP) assay in SW480 cells demonstrating the interaction between endogenous ITGB4 and EZR proteins. (B) Co-immunofluorescence (Co-IF) staining in SW480 cells showing the co-localization of ITGB4 (green) and EZR (red) at the cell membrane. Nuclei were stained with DAPI (blue). (C) Protein-protein interaction network from the STRING database predicting an association between ITGB4 and EZR. (D, E) Verification of EZR overexpression efficiency in SW480 and HCT116 cells by qRT-PCR and Western blot after transfection with pcDNA3.1-EZR. (F, G) qRT-PCR and Western blot analysis showing decreased EZR mRNA and protein levels in SW480 cells after ITGB4 knockdown. (H, I) Western blot and qRT-PCR analysis confirming that EZR overexpression does not rescue the siRNA-mediated knockdown of ITGB4, indicating EZR acts downstream of ITGB4. Data are presented as mean ± SD. *P < 0.05, **P < 0.01, ***P < 0.001.

### ITGB4 promotes CRC progression by regulating EZR to activate the Wnt/β-catenin signaling pathway

To determine the signaling pathway through which the ITGB4/EZR axis functions, we performed Kyoto Encyclopedia of Genes and Genomes (KEGG) pathway enrichment analysis on the DEGs from our RNA-seq data. The analysis revealed a significant enrichment in the Wnt/β-catenin signaling pathway (**Figure 7A**). To validate this, we examined key proteins of the pathway by Western blot. ITGB4 knockdown in HCT116 cells led to a marked decrease in the levels of active β-catenin and its downstream transcriptional targets, c-Myc and Cyclin D1. Crucially, this downregulation was reversed by the ectopic overexpression of EZR (**Figure 7B**). This indicates that ITGB4 activates the Wnt/β-catenin pathway in an EZR-dependent manner. Finally, we conducted functional rescue experiments to confirm that EZR mediates the oncogenic effects of ITGB4. As shown in **Figure 7C-K**, while ITGB4 knockdown suppressed CRC cell proliferation, migration, and invasion, concomitant overexpression of EZR significantly rescued these phenotypes. Taken together, these results demonstrate that ITGB4 promotes CRC progression by regulating EZR, which in turn leads to the activation of the Wnt/β-catenin signaling cascade.

**Figure 7.**
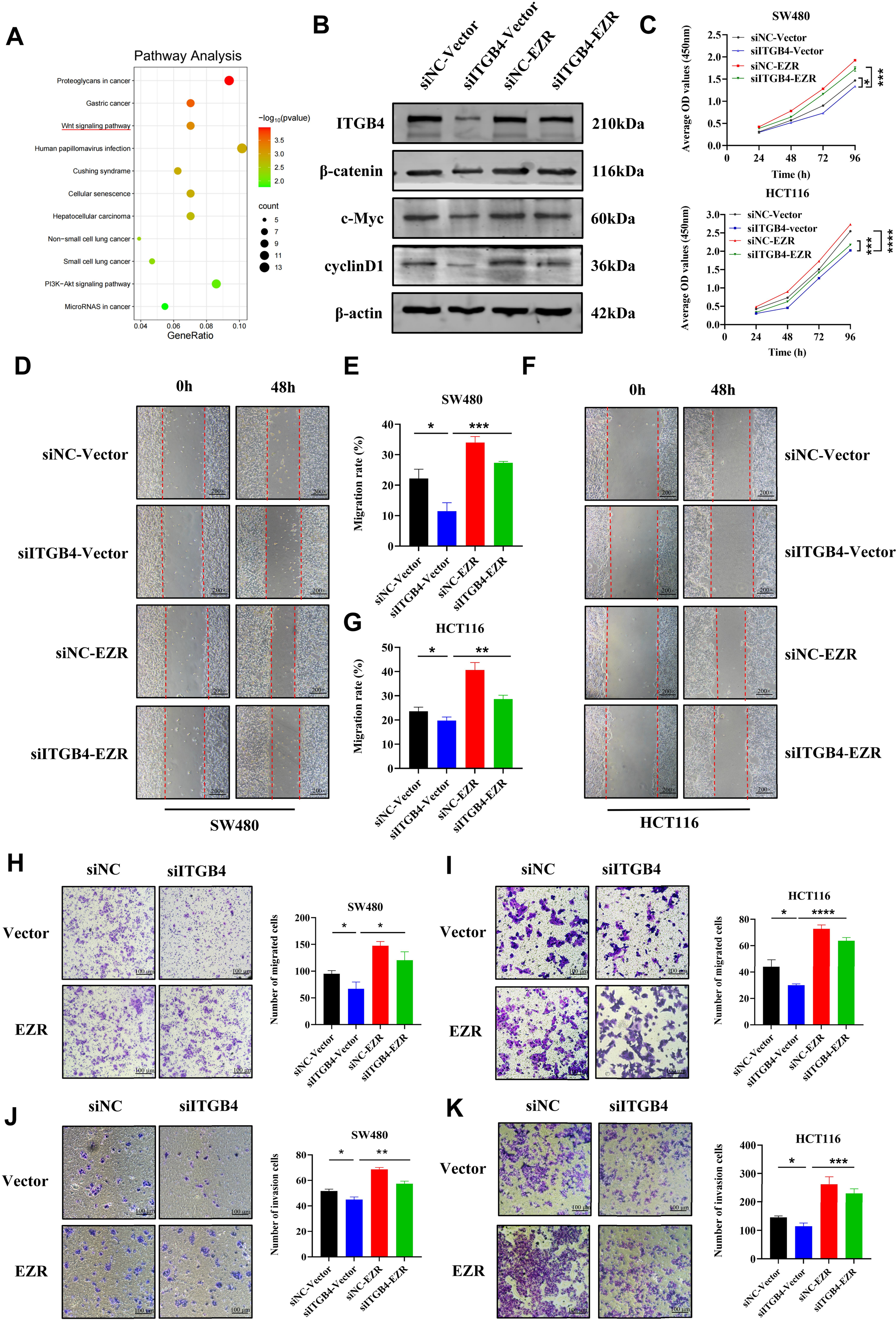
ITGB4 exerts its oncogenic function through the EZR-mediated activation of the Wnt/β-catenin signaling pathway. (A) KEGG pathway enrichment analysis of differentially expressed genes following ITGB4 knockdown, highlighting the Wnt signaling pathway. (B) Western blot analysis showing that ITGB4 knockdown suppresses the expression of β-catenin, c-Myc, and Cyclin D1, and this effect is rescued by EZR overexpression in HCT116 cells. (C) CCK-8 assays demonstrating that EZR overexpression rescues the proliferation defect caused by ITGB4 knockdown in SW480 and HCT116 cells. (D-G) Wound healing assays showing that EZR overexpression restores the migratory capacity inhibited by ITGB4 knockdown (200x magnification). (H, I) Transwell migration assays confirming the rescue of cell migration by EZR overexpression (crystal violet staining, Scale bar = 100 μm). (J, K) Transwell invasion assays showing that EZR overexpression rescues the invasive potential suppressed by ITGB4 knockdown (crystal violet staining, Scale bar = 100 μm). Data are presented as mean ± SD. *P < 0.05, **P < 0.01, ***P < 0.001.

## Discussion

In this study, we have elucidated a novel molecular mechanism underlying the pro-tumorigenic role of ITGB4 in colorectal cancer. We confirmed that ITGB4 is significantly overexpressed in CRC tissues and that its high expression is associated with poor patient prognosis. Through a series of in vitro and in vivo experiments, we demonstrated that ITGB4 is critical for promoting CRC cell proliferation, migration, invasion, and survival. Mechanistically, we identified EZR as a key downstream effector of ITGB4 for the first time. Our data show that ITGB4 interacts with and positively regulates EZR expression, which subsequently activates the Wnt/β-catenin signaling pathway, a well-established driver of CRC. These findings highlight the ITGB4/EZR/Wnt/β-catenin axis as a crucial signaling cascade in CRC progression and suggest its potential as a therapeutic target.

ITGB4, a transmembrane receptor, plays a pivotal role in cell-matrix adhesion and signal transduction [6]. Its aberrant expression has been linked to the progression of various cancers [9–14]. Consistent with these reports and our own previous findings [16], this study solidifies the oncogenic function of ITGB4 in CRC. Our functional experiments clearly show that silencing ITGB4 potently inhibits key malignant phenotypes, including proliferation and invasion, both in cultured cells and in a xenograft tumor model. These results strongly support ITGB4’s role as a driver of CRC and reinforce its value as a potential therapeutic target.

A key innovation of our study is the identification of EZR as a downstream target regulated by ITGB4. Ezrin is a member of the ERM (Ezrin-Radixin-Moesin) protein family that links the plasma membrane to the actin cytoskeleton, thereby regulating cell adhesion, migration, and signaling [19]. EZR is known to be overexpressed in numerous malignancies, including CRC, where its expression correlates with advanced stage, metastasis, and poor prognosis [17, 20–23]. While both ITGB4 and EZR have been independently implicated in cancer, a direct regulatory link between them in the context of CRC has not been previously described. Our study provides compelling evidence for this connection through Co-IP and Co-IF experiments demonstrating their physical interaction and co-localization. The co-localization of ITGB4 and EZR at the plasma membrane is biologically significant. As a transmembrane receptor, ITGB4 requires membrane-proximal effectors to transmit extracellular signals. EZR, serving as a cross-linker between the plasma membrane and the actin cytoskeleton, is ideally positioned to transduce ITGB4-mediated signals to the intracellular machinery. Furthermore, we showed that ITGB4 expression level directly influences EZR mRNA and protein levels, suggesting that ITGB4 may regulate EZR through transcriptional or post-transcriptional mechanisms. While the precise mode of this regulation (e.g., via downstream transcription factors or effects on mRNA stability) warrants further investigation, our results firmly place EZR as a downstream mediator of ITGB4’s oncogenic signaling.

The activation of the Wnt/β-catenin signaling pathway is a hallmark event in the majority of colorectal cancers, driving uncontrolled cell proliferation and survival [24, 25]. The pathway culminates in the nuclear translocation of β-catenin, which acts as a co-activator for TCF/LEF transcription factors to induce the expression of target genes like c-Myc and Cyclin D1 [26]. Our study connects the ITGB4/EZR axis to this critical pathway. We demonstrate that ITGB4 knockdown deactivates the Wnt/β-catenin pathway and that this effect is mediated by EZR, as EZR overexpression could restore pathway activity. This finding is particularly significant as it provides a mechanistic explanation for how an upstream cell adhesion molecule like ITGB4 can influence a core intracellular oncogenic signaling cascade. The ability of EZR to rescue the malignant phenotypes suppressed by ITGB4 knockdown further solidifies the functional importance of this newly identified axis. Furthermore, our data serendipitously revealed that EZR overexpression could partially restore ITGB4 levels. Since β-catenin acts as a prominent transcription factor, it is highly likely that ITGB4 is a transcriptional target of the Wnt/β-catenin pathway. This forms a positive feedback loop (ITGB4/EZR/Wnt/β-catenin/ITGB4) that continuously fuels CRC progression, an intriguing mechanism that warrants dedicated exploration in future studies.

Mechanistically, Ezrin may activate Wnt signaling through several potential pathways. Previous studies in osteosarcoma indicate that Ezrin can physically interact with β-catenin and facilitate its nuclear translocation [27]. Alternatively, Ezrin may disrupt the E-cadherin/β-catenin complex at the cell membrane, releasing a pool of β-catenin that acts as a signaling molecule rather than an adhesion component. Furthermore, Ezrin phosphorylation is a key regulatory event that modulates multiple downstream signaling cascades, including those that converge on β-catenin stability [28]. Our findings in CRC are consistent with these models, suggesting Ezrin acts as a critical signal transducer.

While other integrins, such as Beta 1, have been reported to modulate Wnt signaling via focal adhesion kinase (FAK), the ITGB4-mediated regulation appears distinct. Unlike Beta 1, ITGB4 primarily forms hemidesmosomes. Our data suggests that in cancer cells, ITGB4 signaling is re-wired away from stable adhesion towards pro-migratory signaling via Ezrin, a mechanism that may be unique to the structural properties of the Beta 4 subunit.

Nevertheless, this study has certain limitations. First, while we demonstrated a regulatory relationship, the precise molecular details of how ITGB4 modulates EZR expression require further exploration. Future studies could investigate whether ITGB4 signaling affects transcription factors that bind to the EZR promoter or influences the stability of EZR mRNA. Second, the clinical correlation between ITGB4 and EZR expression should be validated in a larger cohort of CRC patient samples to strengthen its clinical relevance. Additionally, while our in vitro assays strongly support a role for ITGB4 in migration and invasion, the subcutaneous xenograft model used here primarily assesses tumor growth. Future studies utilizing orthotopic implantation or tail vein injection models are necessary to fully validate the metastatic potential of the ITGB4/Ezrin axis in vivo. Despite these limitations, our work provides a robust foundation for future research.

In conclusion, our study identifies ITGB4 as a critical oncogene in colorectal cancer that promotes tumor progression by upregulating its downstream target EZR, leading to the activation of the Wnt/β-catenin signaling pathway. This newly defined ITGB4/EZR/Wnt/β-catenin axis offers novel insights into the molecular pathogenesis of CRC and presents ITGB4 as a promising biomarker for prognosis and a potential target for therapeutic intervention.

## Abbreviation list

CRC: Colorectal Cancer
ITGB4: Integrin Beta 4
EZR: Ezrin
CCK-8: Cell Counting Kit-8
RNA-seq: RNA Sequencing
Co-IP: Co-immunoprecipitation
Co-IF: Co-immunofluorescence
ATCC: American Type Culture Collection
DMEM: Dulbecco’s Modified Eagle’s Medium
FBS: Fetal Bovine Serum
siRNA: small interfering RNA
shRNA: short hairpin RNA
qRT-PCR: Quantitative Real-Time Polymerase Chain Reaction
PVDF: Polyvinylidene Fluoride
HE: Hematoxylin and Eosin
IHC: Immunohistochemistry
DEGs: Differentially Expressed Genes
TCGA: The Cancer Genome Atlas
GEPIA2: Gene Expression Profiling Interactive Analysis 2
OS: Overall Survival
KEGG: Kyoto Encyclopedia of Genes and Genomes
SD: Standard Deviation

## Declarations

### Ethics approval and consent to participate

All animal procedures were approved by the Experimental Animal Care and Use Committee and the Ethics Committee of The First Hospital of Hebei Medical University (Approval No. 20220395). The study was carried out in compliance with the ARRIVE guidelines and all methods were performed in accordance with the relevant guidelines and regulations. This article does not contain any studies with human participants performed by any of the authors.

### Consent for publication

Not applicable.

### Availability of data and materials

The datasets used and/or analysed during the current study are available from the corresponding author on reasonable request.

### Authors’ contributions

WY, JW (Jia Wang), and TL conceived and designed the study. JW (Jing Wang), YS, and MX performed the experiments, analyzed the data, and wrote the manuscript. SH, KL, and JJ provided technical support and study materials. XM and HL contributed to data collection and assembly. All authors read and approved the final manuscript.

## Acknowledgements

Not applicable.

