## Supplementary table 1 for "Integrin beta 4 promotes colorectal cancer progression by upregulating Ezrin and activating the Wnt/β-catenin signaling pathway"

**Supplementary Table 1.** Sequences of siRNAs and shRNAs clones.

| **Name / Description** | **Strand** | **Sequence (5' → 3')** |
| --- | --- | --- |
| **siRNA-ITGB4 sequences** | | |
| ITGB4-Homo-681 | S | GCGACUACACUAUUGGAUUTT |
|  | AS | AAUCCAAUAGUGUAGUCGCTT |
| ITGB4-Homo-822 | S | GUGGAUGAGUUCCGGAAUATT |
|  | AS | UAUUCCGGAACUCAUCCACTT |
| ITGB4-Homo-2454 | S | GCUUUAAGGAAGACCACUATT |
|  | AS | UAGUGGUCUUCCUUAAAGCTT |
| ITGB4-Homo-4033 | S | GCUGCUUAUUGAGAACCUUTT |
|  | AS | AAGGUUCUCAAUAAGCAGCTT |
| **shRNA-ITGB4 sequences** | | |
| shRNA-ITGB4 target sequence | - | GCGACTACACTATTGGATT |
| shDNA template sequence | S | CACCGCGACTACACTATTGGATTTCAAGAGAATCCAATAGTGTAGTCGCTTTTTTG |
|  | A | GATCCAAAAAAGCGACTACACTATTGGATTCTCTTGAAATCCAATAGTGTAGTCGC |
