## Supplementary table 2 for "Integrin beta 4 promotes colorectal cancer progression by upregulating Ezrin and activating the Wnt/β-catenin signaling pathway"

**Supplementary Table 2. Primer sequences used for qRT-PCR.**

| **Gene** | **Forward Primer (5'-3')** | **Reverse Primer (5'-3')** |
| --- | --- | --- |
| ITGB4 | TCTCTCAGAGTGAGCTGGCAG | TTCAGCAGCTGGTACTCCAC |
| EZR | ACCAATCAATGTCCGAGTTACC | GCCGATAGTCTTTACCACCTGA |
| β-actin | CAGCCATGTACGTTGCTATCCAGG | AGGTCCAGACGCAGGATGGCATG |
| EPHA2 | TGGCTCACACACCCGTATG | GTCGCCAGACATCACGTTG |
| ITGA6 | ATGCACGCGGATCGAGTTT | TTCCTGCTTCGTATTAACATGCT |
| CXADR | GTGCTCCTGTGCGGAGTAG | ATGGCAGATAGGCAGTTTCCC |
| GPRC5C | CCTGTACTACAACCTGTGTGAC | TGAGCACAAACGTGGTGACA |
